# T cell receptor repertoire signatures associated with COVID-19 severity

**DOI:** 10.1101/2021.11.30.470640

**Authors:** Jonathan J. Park, Kyoung A V. Lee, Stanley Z. Lam, Sidi Chen

**Affiliations:** Department of Genetics, Yale University School of Medicine, New Haven, CT, USA; System Biology Institute, Yale University, West Haven, CT, USA; Center for Cancer Systems Biology, Yale University, West Haven, CT, USA; M.D.-Ph.D. Program, Yale University, West Haven, CT, USA; Molecular Cell Biology, Genetics, and Development Program, Yale University, New Haven, CT, USA; Department of Biostatistics, Yale University School of Public Health, New Haven, CT, USA; Immunobiology Program, Yale University, New Haven, CT, USA; Yale Comprehensive Cancer Center, Yale University School of Medicine, New Haven, CT, USA; Yale Stem Cell Center, Yale University School of Medicine, New Haven, CT, USA; Yale Center for Biomedical Data Science, Yale University School of Medicine, New Haven, CT, USA

## Abstract

T cell receptor (TCR) repertoires are critical for antiviral immunity. Determining the TCR repertoires composition, diversity, and dynamics and how they change during viral infection can inform the molecular specificity of viral infection such as SARS-CoV-2. To determine signatures associated with COVID-19 disease severity, here we performed a large-scale analysis of over 4.7 billion sequences across 2,130 TCR repertoires from COVID-19 patients and healthy donors. TCR repertoire analyses from these data identified and characterized convergent COVID-19 associated CDR3 gene usages, specificity groups, and sequence patterns. T cell clonal expansion was found to be associated with upregulation of T cell effector function, TCR signaling, NF-kB signaling, and Interferon-gamma signaling pathways. Machine learning approaches accurately predicted disease severity for patients based on TCR sequence features, with certain high-power models reaching near-perfect AUROC scores across various predictor permutations. These analyses provided an integrative, systems immunology view of T cell adaptive immune responses to COVID-19.

## Introduction

Much of the ongoing COVID-19 vaccination strategies have focused on targeting B cells for eliciting neutralizing antibodies (nAbs) against SARS-CoV-2 ^1,2^. However, SARS-CoV-2 nAb levels after infection or vaccination have been found to decrease over time ^3^, and recently emerging variants of concern (VOC) have been associated with antibody escape ^4^. Strategies that solely focus on nAbs may not be sufficient for managing the pandemic in the long term. There has therefore been increasing interest in studying the role of T cell immunity in the response to COVID-19 infection ^5,6^.

Functional T cell responses are crucial for control and clearance of many respiratory viral infections ^7^, including for SARS-CoV and MERS-CoV ^8,9^. Studies from transgenic mouse models suggest that T cells are also important for disease resolution after infection with SARS-CoV-2 ^10^, and SARS-CoV-2-specific CD4 and CD8 T cells have been associated with milder disease in human patients ^11^, suggesting roles for coordinated adaptive immune responses in protective immunity against COVID-19. T cells contribute to viral control through numerous mechanisms, including supporting the generation of antibody-producing plasma cells (T follicular helper cells), production of effector cytokines such as IFN-gamma and TNF, and cytotoxicity against infected cells. Generation of memory T cells can provide life-long protection against pathogens ^12^, and a recent study showed that SARS-CoV-2-specific memory T cell responses were sustained for 10 months in COVID-19 convalescent patients ^13^. Moreover, there is mounting evidence that SARS-CoV-2 VOCs rarely escape T reactivity ^14^, perhaps in part due to a wider distribution of T cell epitopes across the entire viral proteome unlike nAb target limitation to the viral surface. Due to the importance of T cells in long-term and broad immune reactivity, there has been an increase in diverse vaccine strategies to expand targets beyond the spike protein and induce T cell responses ^5^.

T cells recognize viral antigens presented on major histocompatibility complex (MHC) molecules through an enormously diverse assembly of T cell receptors (TCRs) ^15^. Ligation of the TCR by peptide-loaded MHC molecules leads to T cell activation and clonal expansion, causing a shift in repertoire specificity towards the antigen. TCR repertoires therefore represent a functional signature of the adaptive immune response. The development of high-throughput DNA sequencing methods has enabled highly quantitative investigation into the diversity and composition of immune repertoires ^16^; for example, one study used TCR sequencing (TCR-seq) on samples from T-lineage acute lymphoblastic leukemia/lymphoma patients to reveal the receptor profiles of clonal T lymphoblast populations and then further to develop a clinical assay for diagnosis of minimal residual disease ^17^. Other studies have used tracking of TCR repertoires in cancer patients over time to identify correlations between clonal dynamics and clinical features such as immunotherapy treatment response ^18,19^. TCR-seq data has enormous potential for gaining quantitative insight into the patterns of adaptive immune responses, which has been particularly well demonstrated in studies for cancer immunology.

We sought to develop an integrative, systems immunology approach for investigating TCR repertoires from COVID-19 patients to help decode patterns of the adaptive immune response during SARS-CoV-2 infection. While there have been some preliminary studies on different aspects of TCR-seq analysis for COVID-19 ^20–23^, there have been limited studies that incorporate motif-based analysis, transcriptomics, and machine learning in a large-scale, comprehensive investigation into the immune responses during disease course of varying severity. We anticipate that our approach here can provide sets of COVID-19 associated sequences and motifs that may help guide development of prognostic and diagnostic markers and potentially help design therapeutic interventions that better harness the power of T cell immunity.

## Results

### TCR repertoires from COVID-19 patients and healthy donors reveal trends in CDR3 gene usage and diversity

To determine if there were any global patterns that distinguish the immune repertoires of COVID-19 patients, we systematically compiled and analyzed TCR-seq samples (total n = 2130) from COVID-19 patients and healthy donors (**Figure 1A**). TCR repertoire data was obtained from studies by Adaptive Biotechnologies (AB, n = 1574), ISB-Swedish COVID-19 Biobanking Unit (ISB-S, n = 266), PLA General Hospital (PLAGH, n = 20), and Wuhan Hankou Hospital (WHH, n = 15), and then uniformly processed for downstream analysis (see **Methods**). Clonality analyses revealed that COVID-19 patient samples from the ISB-S CD4, ISB-S CD8, and WHH datasets had significantly fewer total unique clonotypes compared to healthy donor controls (**Figure S1A**). Moreover, repertoire diversity metrics including Chao1 estimators (measure of species richness), Gini-Simpson indices (probability of interspecific encounter), and inverse Simpson indices were significantly decreased for COVID-19 samples compared to healthy donor samples, notably for the AB, ISB-S CD4, and ISB-S CD8 datasets (**Figures 1B-C, S1B**). The decrease in clonal diversity measures is consistent with the increase in the relative abundance of the top clonotypes in the repertoire space for COVID-19 samples (**Figures 1F, S3D**), which suggests expansion of a small number of functional clones after antigen exposure. These results together reveal global shifts in immune repertoire clonality and diversity in patients with COVID-19 compared to healthy donors.

**Figure 1.**
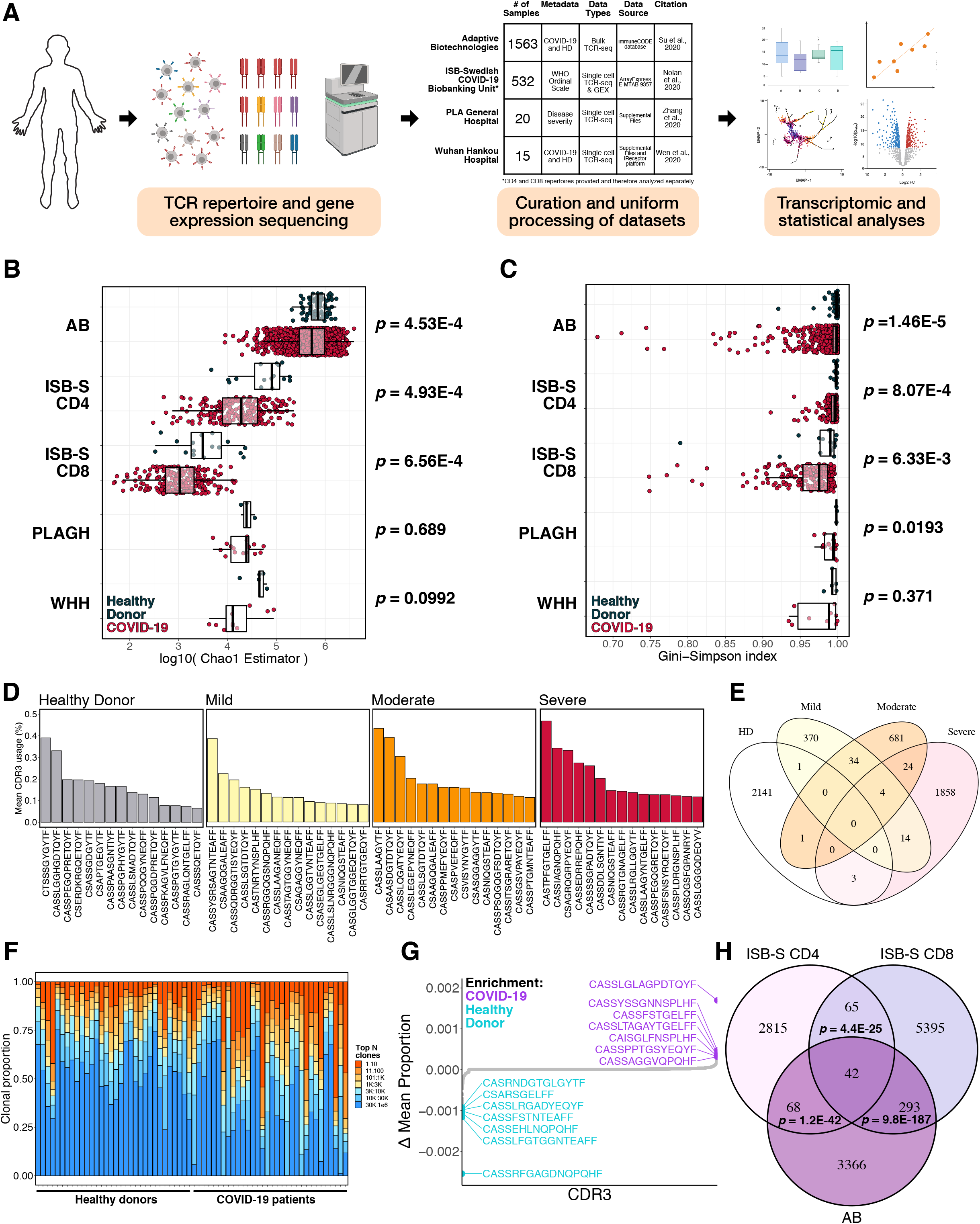
Analysis of TCR repertoires from COVID-19 patients and healthy donors reveal trends in CDR3 gene usage and diversity. **(A)** Schematic detailing curation and analysis of TCR repertoire datasets from healthy donors and COVID-19 patients. Sequencing data was obtained from Adaptive Biotechnologies (AB, n = 1574), ISB-Swedish COVID-19 Biobanking Unit (ISB-S, n = 266, CD4 and CD8 repertoires), PLA General Hospital (PLAGH, n = 20), and Wuhan Hankou Hospital (WHH, n = 15). **(B)** Boxplot of Chao1 indices for COVID-19 patients and healthy donors for each repertoire dataset. P-values were obtained using the two-sided Wilcoxon rank sum test. **(C)** Boxplot of Gini-Simpson indices for COVID-19 patients and healthy donors for each repertoire dataset. P-values were obtained using the two-sided Wilcoxon rank sum test. **(D)** Bar plots showing the top 15 mean CDR3 usages for patients in the ISB-S CD4 dataset grouped by disease severity (healthy donor = 16, mild = 108, moderate = 93, severe = 49). **(E)** Venn diagram showing overlap of top mean CDR3 usages (proportion threshold = 0.0001) for patients in the ISB-S CD4 dataset grouped by disease severity. **(F)** Bar plot depicting relative abundance for groups of top clonotypes for sampled repertoires (healthy donors = 32, COVID-19 = 32) from AB dataset. **(G)** Dotted waterfall plot of CDR3 gene usage differentials between COVID-19 patients and healthy donors (delta mean proportion) in AB dataset. Purple dots are CDR3 sequences enriched in COVID-19; light blue dots are CDR3 sequences enriched in healthy donors; grey dots are all other CDR3 sequences. **(H)** Venn diagram showing overlap of COVID-19 enriched CDR3 sequences for patients in the ISB-S CD4, ISB-S CD8, and AB datasets (thresholds 0.0001 for ISB-S samples, 0.00001 for AB samples). P-values for overlap significance calculated using hypergeometric test.

To determine the specific gene usage preferences and dynamics in COVID-19 patients, we performed comparative analyses of V(D)J gene and complementarity-determining region 3 (CDR3) gene usages for the AB and ISB-S datasets. While we observed some significant selective V and J gene usage differences in the AB dataset (**Figures S1D-E**), fewer differences were found for the ISB-S CD4 and CD8 datasets when comparing samples from different disease severities to those from healthy donors (**Figures S2A-D**). Moreover, there were no differences in clonotype frequencies by CDR3 length across the datasets (**Figures S1C**). By comparison, the top CDR3 sequences were different across conditions for both the AB and ISB-S datasets (**Figures 1D, S3A, S3B**). In order to identify COVID-19 associated CDR3 sequences that are conserved across disease conditions and datasets, we performed a series of set analyses using sequences above a proportion threshold (0.0001 for ISB-S samples, 0.00001 for AB samples) for each condition. We found that CDR3 sequences enriched in the mild, moderate, and severe disease condition samples from the COVID-19 patients had considerable overlap while having limited overlap with healthy donor samples for both the ISB-S CD4 (**Figure 1E**) and ISB-S CD8 (**Figure S3B**) datasets. Moreover, we observe 42 conserved CDR3 sequences when comparing the union set of disease-associated CDR3 sequences for ISB-S CD4 samples, the union set of disease-associated CDR3 sequences for ISB-S CD8 samples, and COVID-19 CDR3 sequences for the AB samples (**Figure 1H**). In order to determine enriched CDR3 sequences for each dataset and disease conditions, we plotted the difference in mean CDR3 proportions between samples of interest and healthy donors (**Figures 1G, S3E-J**). Although the identified sequences may not be definitively specific, we provide here a set of systematically processed COVID-19 associated convergent and enriched CDR3 gene usages.

### K-mer and motif analyses reveal patterns associated with disease conditions

Sequence convergence of immune repertoires can be also occur at the level of motifs, or sequence substrings, in addition to that of clones. One approach to decomposing CDR3 sequences into motifs is by using overlapping k-mers, or amino acid sequences of length k, which provide a functional representation of the repertoires with increased compatibility for statistical analyses and machine learning methods ^24^. We created 3-mer, 4-mer, 5-mer, and 6-mer frequency matrix representations of ISB-S CD4 and CD8 datasets and performed principal components analysis to see whether samples cluster by disease severity (**Figures 2A, 2C, S4A-F**). We found that while the majority of samples clustered together, a number of mild and moderate samples were separated from the main cluster across all analysis permutations, while severe samples are generally associated with the main cluster of samples including healthy donors. These results are consistent emerging data that patients with severe COVID-19 have substantial immune dysregulation in comparison to those with less severe disease. Studies have shown that T cell polyfunctionality is increased in patients with moderate disease but reduced in those with severe disease ^25^, and there have been proposed models of TCR clonality whereby the response in mild disease includes detection of dominant clones while response in severe disease do not ^26^. Moreover, heatmaps of 3-mer abundances reveal some shared motifs between mild and moderate samples such as YNE, NEQ, EQF, and QFF for repertoires sampled from the ISB-S CD4 dataset and TEA, EAF, and AFF for repertoires sampled from the ISB-S CD8 dataset (**Figures 2B, 2D**). These results in aggregate suggest that there are sequence features that distinguish COVID-19 TCR repertoires from healthy donors to various degrees based on disease condition.

**Figure 2.**
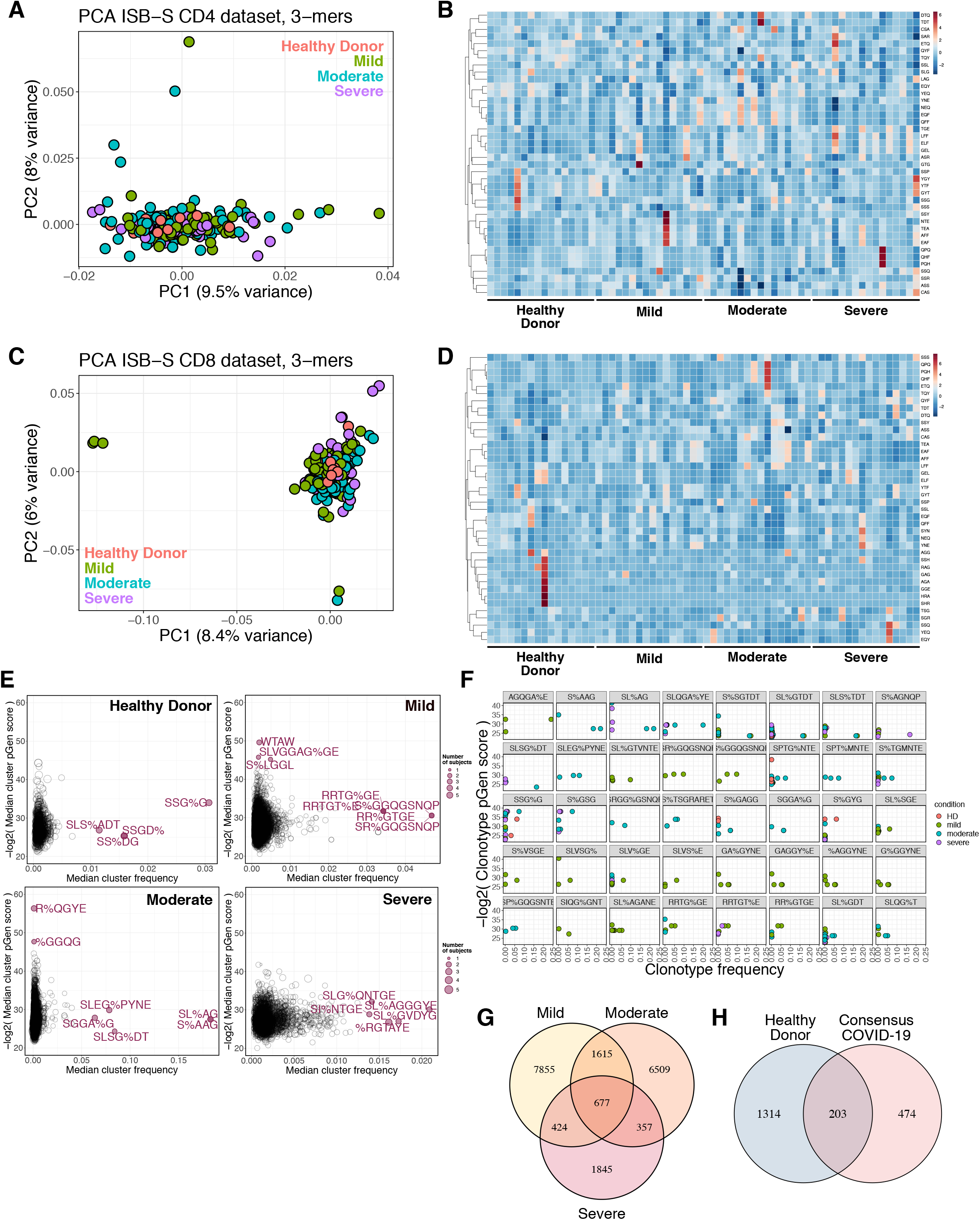
K-mer and motif analyses reveal patterns associated with disease condition. **(A)** Principal components analysis of 3-mer representations of TCR repertoires from the ISB-S CD4 dataset (n = 266). **(B)** Heatmaps of 3-mer abundances of repertoires sampled from the ISB-S CD4 dataset by disease condition (healthy donor = 16, mild = 16, moderate = 16, severe = 16). **(C)** Principal components analysis of 3-mer representations of TCR repertoires from the ISB-S CD8 dataset (healthy donor = 16, mild = 108, moderate = 93, severe = 49). **(D)** Heatmaps of 3-mer abundances of repertoires sampled from the ISB-S CD8 dataset by disease condition (n = 266). **(E)** Median frequency and pGen scores of COVID-19 and healthy donor associated T cell clusters from GLIPH2 analysis of the ISB-S CD4 dataset, grouped by disease condition. **(F)** Detailed view of frequencies and pGen scores of specific clonotypes associated with high frequency T cell clusters from CD4 dataset. Clonotypes are colored by patient disease condition. **(G)** Venn diagram showing overlap of COVID-19 associated T cell clusters for patients in the ISB-S CD4 dataset grouped by disease condition. **(H)** Venn diagram showing overlap between consensus COVID-19 associated T cell clusters (taken from intersection of disease conditions) and healthy donors for repertoires in the ISB-S CD4 dataset.

Recent sequence similarity approaches have been developed to determine TCR specificity clusters for motif-based prediction of antigen specificity and identification of key conserved residues that drive TCR recognition ^27–29^. We used the Grouping of Lymphocyte Interactions by Paratope Hotspots version 2 (GLIPH2) algorithm ^29^ to cluster the TCR sequences based on predicted antigen specificity for significant motifs associated with different disease conditions in the ISB-S datasets. We also used the Optimized Likelihood estimate of immunoGlobulin Amino-acid sequences (OLGA) algorithm ^30^ to calculate the generation probability (pGen) of the clonotypes contained in the clusters identified from the GLIPH2 analysis. Low pGen clonotypes are considered private and not shared widely in the population, while high pGen clonotypes are considered public and shared in a large proportion of the population due to convergent recombination ^20,31^. We found that the mild and moderate disease conditions had both relatively lower pGen scores and higher median frequency clusters compared to the severe disease and healthy donor conditions for both the ISB-S CD4 and CD8 datasets (**Figures 2E, S4G**). Visualization of individual clusters revealed that the mild and moderate disease conditions had clonotypes with the highest proportional representation, including motifs AGQGA%E, S%AAG, SL%AG, and SLQGA%YE (the % character corresponds to a wildcard amino acid) for the ISB-S CD4 dataset (**Figure 2F**) and motifs SEG%NTDT, SLDSGGA%E, SL%SGGANE, SLAA% for the ISB-S CD8 dataset (**Figure S4H**). In order to identify clusters that were exclusive to COVID-19 patients in the ISB-S CD4 dataset, we performed set analysis and found 677 clusters in the intersection of the disease conditions, 474 of which were exclusive and had no overlap with the healthy donor clusters (**Figures 2G-H**). For the ISB-S CD8 dataset, we found 51 consensus clusters, 35 of which were exclusive (**Figures S4I-J**). We provide here all identified clusters and motifs with associated CDR3 sequences, V gene usage, and J gene usages, along with clonotype pGen scores and the identified COVID-19 associated clusters.

### Transcriptional signatures of clonal expansion and associations with disease severity

In order to investigate the relationship between the enriched clonotypes and their transcriptomes, we performed dimensionality reduction on 137,075 CD4 T cell single cell RNA sequencing samples that had CDR3 sequences associated with identified GLIPH2 clusters. The transcriptomes were projected to a two dimensional space by uniform manifold approximation and projection (UMAP) (**Figure S5A**). Clustering was performed using the Louvain algorithm, revealing 12 clusters with differentially expressed gene signatures (**Figures 3A, S5C**). We found that cluster 6 contained cells with high degrees of clonality (**Figures 3A, S5B**), suggesting phenotypic correlates of clonal expansion. Comparison with the top enriched motifs found from the GLIPH2 analysis, including AGQGA%E, S%AAG, SL%AG, SLQGA%YE, S%SGTDT, SL%GTDT, SLS%TDT, and S%AGNQP revealed high density of clusters in cluster 6 (**Figures 3C, 2F**). Moreover, we found a correlation between clonotype expansion and disease severity, with cells from COVID-19 patients exhibiting the highest density in effector phenotype associated cluster 6, while healthy donor cells exhibiting density in the naïve phenotype associated clusters (**Figure 3B**). We also found a higher association of lower pGen score, or private, clonotypes with cluster 6 compared to the high pGen score clonotypes (**Figure 3D**), suggesting that these clones may be specific. However, comparison of the proportion of cells for each disease condition in cluster 6 with healthy donors revealed statistically significant cell proportion increases only for the moderate condition (**Figure 3E**), despite increasing trends for all conditions. Altogether, these results demonstrate relationships between clonal expansion, disease severity, and cell phenotype, which can be extended to subsequence motifs.

**Figure 3.**
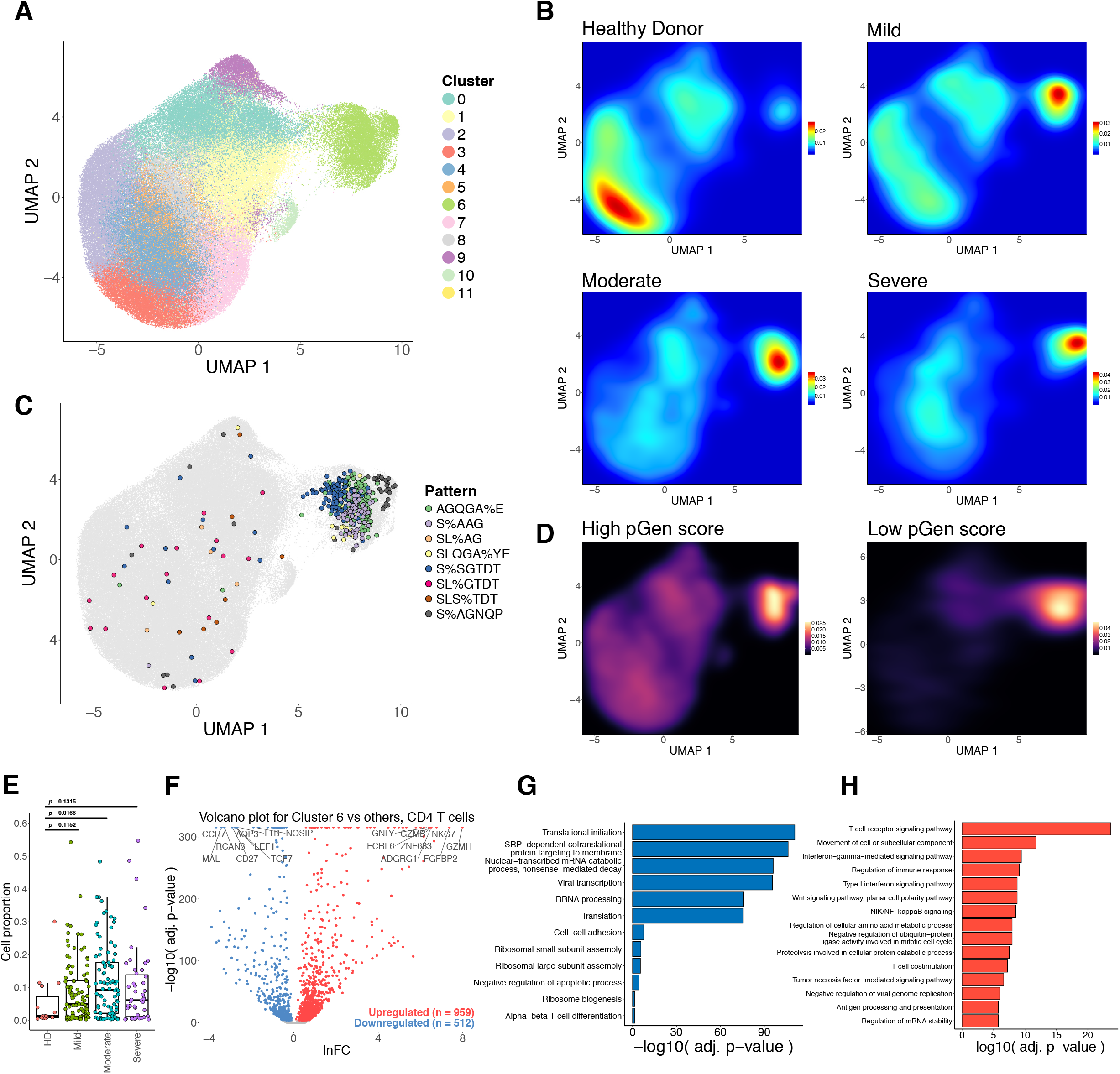
Single cell transcriptional signatures of clonal expansion of CD4 T cells. **(A)** UMAP visualization of 137,075 CD4 T cell single cell transcriptomes from the ISB-S CD4 dataset pooled across samples and conditions. 12 clusters identified using the Louvain algorithm. **(B)** Two dimensional density plot of cells from each disease condition (healthy donor, mild, moderate, severe) by UMAP coordinates. Red represents areas of high density of cells of a given condition; blue represents areas of low density. **(C)** UMAP visualization with cells labelled by top eight most frequent CD4 TCR clusters identified by the GLIPH2 analysis. **(D)** Two dimensional density plot of cells with high or low pGen score clonotypes by UMAP coordinates. Yellow represents areas of high density of cells; black represents areas of low density. **(E)** Boxplots of clonally expanded cell proportions (in cluster 6) for each disease condition (cell count healthy donor = 544, mild = 3,568, moderate = 5,012, severe = 2,336). Comparison between groups performed with two-sided Wilcoxon rank-sum test. **(F)** Volcano plot of differentially expressed genes between clonally expanded cells and all other cells in the ISB-S CD4 dataset (Cluster 6 cells = 5,000, all other cells = 5,000). Differential gene expression was performed with Seurat using the two-sided Wilcoxon rank-sum test; the Bonferroni corrected adjusted p-values and log fold-change of the average expression were used for visualization. **(G)** Bar plot of biological processes (BP) pathway terms associated with downregulated genes (clonally expanded cells vs all other cells, q-value < 1e-4) by DAVID analysis. **(H)** Bar plot of biological processes (BP) pathway terms associated with upregulated genes (clonally expanded cells vs all other cells, q-value < 1e-4) by DAVID analysis.

We extended this analysis to the CD8 dataset to see if the associations between clonal expansion and disease severity are maintained. UMAP projection of 70,237 CD8 T cell single cell transcriptomes and clustering revealed 15 clusters with differentially expressed gene signatures (**Figures 4A, S6A, S6C**). As with the CD4 dataset, we found clustering of cells with high degrees of clonality, distributed here across the clusters 0, 2, 3, 5, 7, 9, 10, 13, and 14 (grouped together as “Expanded” for further analysis) (**Figures 4A, S6B**). We also found high density of top enriched GLIPH2 motifs in the Expanded group, including SEG%NTDT, SLDSGGA%E, SL%SGGANE, SLAA%, SQT%STDT, SP%SGSYE, SPGT%GYNE, and S%RQGAGGE (**Figures 4C, S4H**). We observe a relatively higher density of cells from COVID-19 disease conditions in the Expanded group as compared to the healthy donors (**Figures 4B**), with low density of disease-associated cells in the non-Expanded clusters. Likewise, we found a more exclusive association between lower pGen score clonotypes and the Expanded group, particularly cluster 9 (**Figures 4D**). Comparison of the proportion of cells for each disease condition in the Expanded group with healthy donors reveal statistically significant cell proportion increases for all conditions (**Figure 4E**). These results highlight the relationship between clonal expansion and disease severity comparable with those from the CD4 dataset.

**Figure 4.**
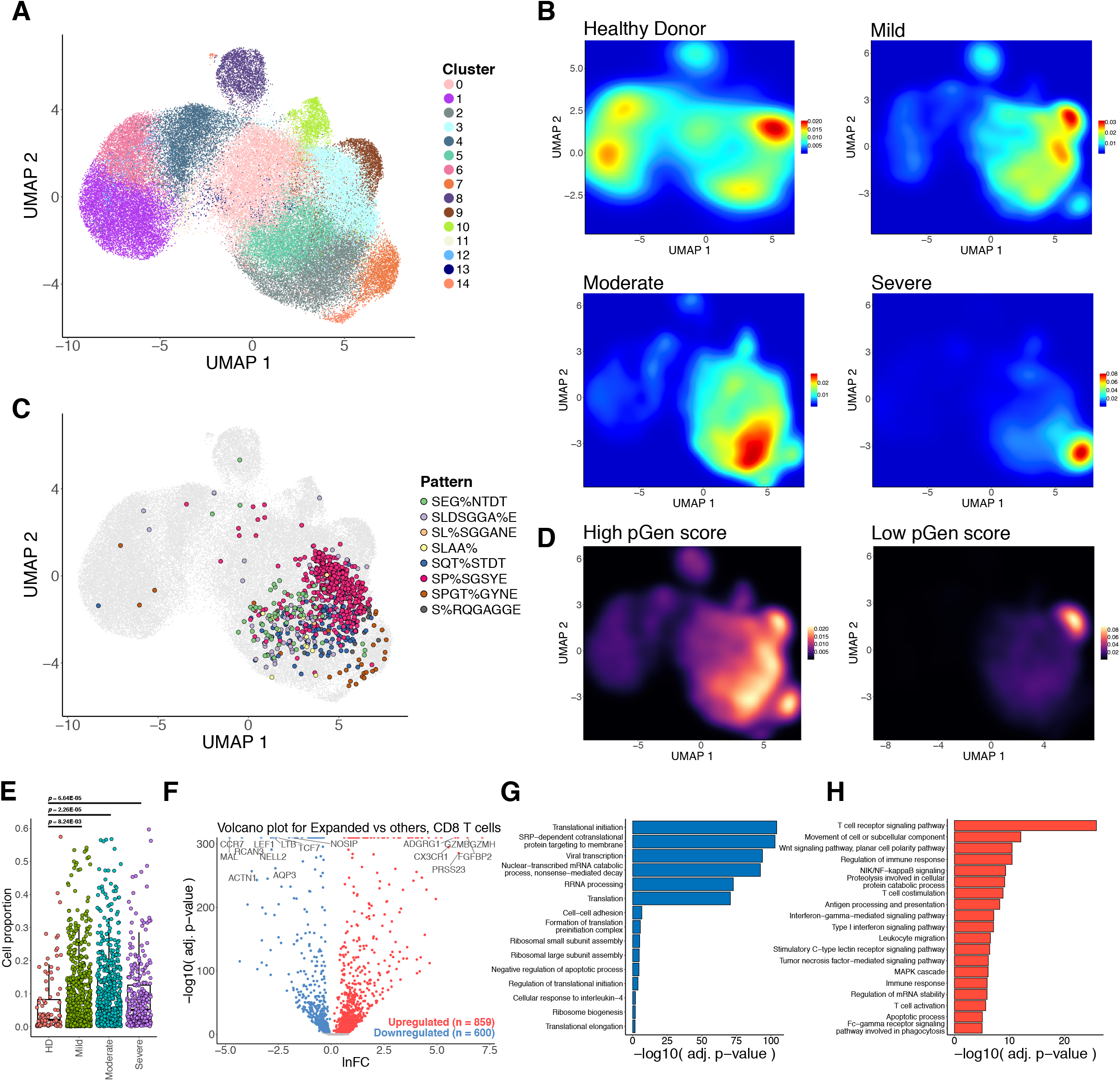
Single cell transcriptional signatures of clonal expansion of CD8 T cells. **(A)** UMAP visualization of 70,237 CD8 T cell single cell transcriptomes from the ISB-S CD8 dataset pooled across samples and conditions. 15 clusters identified using the Louvain algorithm. **(B)** Two dimensional density plot of cells from each disease condition (healthy donor, mild, moderate, severe) by UMAP coordinates. Red represents areas of high density of cells of a given condition; blue represents areas of low density. **(C)** UMAP visualization with cells labelled by top eight most frequent CD8 TCR clusters identified by the GLIPH2 analysis. **(D)** Two dimensional density plot of cells with high or low pGen score clonotypes by UMAP coordinates. Yellow represents areas of high density of cells; black represents areas of low density. **(E)** Boxplots of clonally expanded cell proportions (in Expanded group) for each disease condition (cell count healthy donor = 2,579, mild = 18,622, moderate = 15,743, severe = 7,159). Comparison between groups performed with two-sided Wilcoxon rank-sum test. **(F)** Volcano plot of differentially expressed genes between clonally expanded cells and all other cells in the ISB-S CD8 dataset (Expanded group cells = 5,000, all other cells = 5,000). Differential gene expression was performed with Seurat using the two-sided Wilcoxon rank-sum test; the Bonferroni corrected adjusted p-values and log fold-change of the average expression were used for visualization. **(G)** Bar plot of biological processes (BP) pathway terms associated with downregulated genes (clonally expanded cells vs all other cells, q-value < 1e-4) by DAVID analysis. **(H)** Bar plot of biological processes (BP) pathway terms associated with upregulated genes (clonally expanded cells vs all other cells, q-value < 1e-4) by DAVID analysis.

To investigate the gene expression changes that occur with clonal expansion, we performed differential expression (DEX) analysis between cluster 6 cells versus all other cells for the CD4 dataset (**Figure 3F**) and the Expanded group cells versus all other cells for the CD8 dataset (**Figure 4F**). Using a threshold of q-value < 1e-4, we found 512 downregulated genes and 959 upregulated genes for the CD4 T cell DEX, as well as 600 downregulated genes and 859 upregulated genes for the CD8 T cell DEX. Volcano plots for both T cell types revealed upregulation of cytotoxicity associated transcripts such as granzymes and granulysin and downregulation of naïve phenotype associated markers such as TCF7 and LEF1. Comparison of UMAPs of the individual subpopulation phenotype markers also showed correlation between cluster 6 or the Expanded group clusters and effector-related markers such as GZMA, PRF1, NKG7, and GNLY, with downregulation of naïve-related markers such as TCF7 and LEF1 (**Figures S5D, S6D**). Functional gene annotation analysis with DAVID ^32^ revealed enriched pathways terms such as TCR signaling pathway, regulation of immune response, NF-kB signaling, IFN-gamma mediated signaling, and TNF-mediated signaling pathways were upregulated in clonally expanded clusters (**Figures 3H, 4H**) while terms such as translational initiation, viral transcription, translation, and ribosomal subunit assembly were downregulated (**Figures 3G, 4G**) for both CD4 and CD8 differential expression analyses. We therefore find that clonally expanded CDR3 sequences and motifs are highly associated with effector T cell phenotypes at both the individual gene and functional pathway levels, while downregulating a number of mRNA processing related programs.

### Machine learning models for disease severity

To determine whether the constitutive sequence motifs in the CDR3 sequence of the TCR contain sufficient information to be predictive of disease severity in COVID-19 infection, we trained several classical supervised machine learning (ML) algorithms on the repertoires from the ISB-S CD4 and CD8 datasets. We implemented Random Forests (RF), Support Vector Machines (SVM), Naïve Bayes (NB), Gradient Boosting Classifiers (GBC), and K-Nearest Neighbors (KNN) on frequency matrices of overlapping 3-mer or 6-mer amino acids adapted from the TCR repertoires. ML models were trained as binary classification tasks to predict mild, moderate, or severe COVID-19 TCR repertoires from healthy donor repertoires for either CD4 or CD8 ISB-S datasets. Training and testing partitions were created as five randomly sampled folds, with models trained on 80% of the data and tested on the remaining 20%. This process was repeated 100 times for each fold (500 repetitions per model per classification permutation) for statistical power. We found that RFs, GBCs and SVMs had particularly strong classification performance across the board, with average AUROCs greater than 0.90 for all permutations, with certain predictors approaching perfect score (AUROCs = 0.99 – 1.00) (**Figure 5A, 7A**), as compared to NBs or KNNs. Notably, the ML models had higher performance for classifying mild and moderate repertoires than severe repertoire regardless of k-mer or T cell type. This is consistent with the increased separation of the mild and moderate repertoires observed in the PCA analysis. Overall, these results demonstrate that ML-based methods are capable of identifying samples with high performance from COVID-19 patients of varying severity based on CDR3 sequences features, particularly for mild and moderate disease conditions.

**Figure 5.**
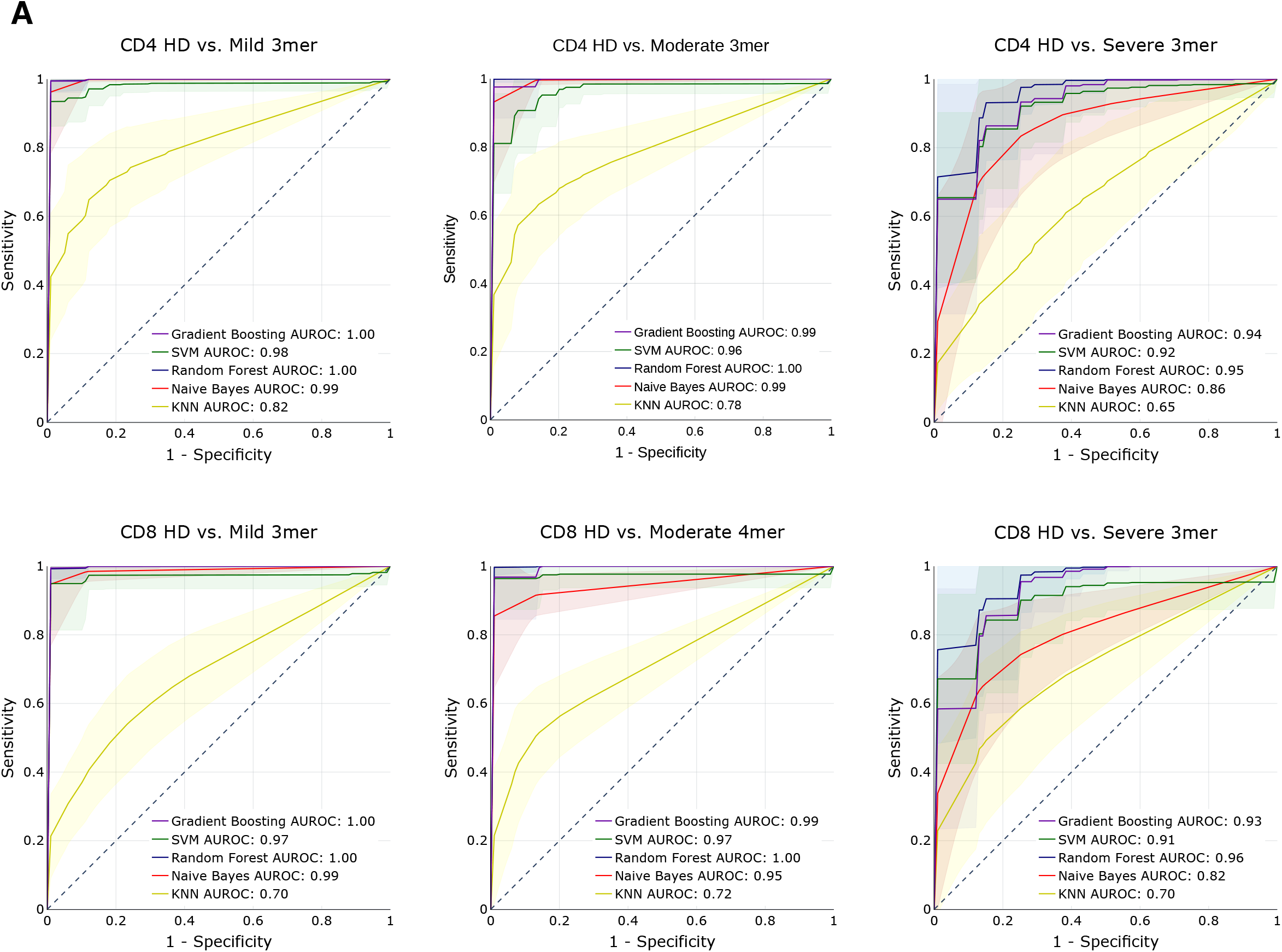
Predictive performance of machine learning models for disease severity. **(A)** AUROC curves for five machine learning models (gradient boosting trees, support vector machines, random forests, Naïve Bayes, and k-nearest neighbors) using 3-mer representations of TCR repertoire data. Models were trained to predict disease severity (mild, moderate, severe) vs healthy donors for CD4 (top row) and CD8 (bottom row) samples. Training and evaluation was performed using 100 repetitions of 5-fold cross-validations per model, average performance +/− 1 standard deviation shown on individual plots.

## Discussion

T cells are increasingly being recognized as key mediators of viral clearance and host protection in COVID-19, and are subjects of active investigation ^33–35^. However, the rules governing SARS-CoV-2 responsive T cell specificity are still incompletely understood. We provide here a comprehensive, systems immunology approach to analyzing COVID-19 TCR repertoires to discover these rules in an unbiased and systematic manner. By uniformly processing immune sequencing data from multiple cohorts with TCR-seq data, we found that antigen exposure during the course of COVID-19 significantly decreased the diversity of repertoires and reshaped clonal representation. We identified and characterized enriched CDR3 sequences, k-mer motifs, and patterns associated with disease severity, and found convergent CDR3 gene usages and clusters that have potential for clonal tracking studies. Comparison of COVID-19 associated motifs and single T cell transcriptomes revealed associations between clonal expansion, disease severity, and cell phenotypes such as effector T cell function. Finally, we established several ML methods for predicting disease severity from TCR repertoires, demonstrating high performance for several models and the potential of using ML for prognostication in COVID-19 patients.

Recent studies have started to report on the differences between T cell responses during mild and more severe COVID-19 disease course. Notably, severe COVID-19, albeit having increases in activated effector cell populations as seen with other disease severities, is associated with lymphopenia and profound functional impairment of CD4 and CD8 T cells ^26,36–40^. These results are consistent with our PCA and motif analyses, we observe a stronger signal for the mild and moderate disease repertoires distinguishing them from healthy donors as compared to severe disease repertoires. Moreover, our ML-based methods had higher performance for predicting mild and moderate repertoires, further demonstrating that CDR3 sequence and subsequence features for these disease conditions have higher discriminative capacity than for severe conditions. Nevertheless, all of the disease conditions were well differentiated from healthy donors across all analyses suggesting that consensus disease-associated features can be identified, including correlations between clonal expansion in the setting of COVID-19 with effector T cell functions at the transcriptomic level.

To our knowledge this is the largest scale investigation into TCR specificity groups for COVID-19 to date, spanning 4,730,447,888 clones across 2,130 repertoires. Though many studies have sought to identify factors predictive of COVID-19 clinical course and outcomes ^41^, few have leveraged TCR-seq data and adaptive immune profiles to their full capacity. We provide high confidence convergent COVID-19 associated signatures with potential prognostic value, including successful implementation of machine learning models for predicting disease severity. The use of next-generation sequencing of immune repertoires provides deeper and more quantitative understanding of the adaptive immune response to COVID-19, and may guide patient risk stratification, vaccine design, and improved clinical management.

## Acknowledgments

We thank various members from Chen lab for discussions. We thank staffs from Yale HPC for technical support. We thank various support from Department of Genetics; Institutes of Systems Biology and Cancer Biology; Dean’s Office of Yale School of Medicine and the Office of Vice Provost for Research.

## Author contributions

JJP and SC conceived and designed the study. JJP developed the analysis approach, performed data analyses, and created the figures. KVL performed data analyses and established machine learning pipelines. SZL performed pre-processing of datasets. JJP and SC prepared the manuscript with input from KVL and SZL. SC supervised the work.

## Methods

### Sequence data collection

TCR repertoire data was obtained from datasets published by Adaptive Biotechnologies, ISB-Swedish COVID-19 Biobanking Unit ^25^, Fifth Medical Center of PLA General Hospital ^21^, and Wuhan Hankou Hospital China ^22^. For COVID-19 patients sequenced with Adaptive Biotechnologies immunoSEQ assays, TCR-seq data were obtained from the ImmuneCODE database at https://doi.org/10.21417/ADPT2020COVID; for healthy donor patients, TCR-seq data was obtained at https://doi.org/10.21417/ADPT2020V4CD. Single cell TCR-seq and gene expression (GEX) data for CD4+ and CD8+ T cell repertoires from COVID-19 patients and healthy donors from the ISB-Swedish COVID-19 Biobanking Unit ^25^ was obtained from the ArrayExpress database ^42^ (http://www.ebi.ac.uk/arrayexpress) using the accession number E-MTAB-9357. Single cell TCR-seq data from COVID-19 patients and healthy donors were also obtained from the Fifth Medical Center of PLA General Hospital, accessed through the supplementary tables of the associated publication ^21^; and Wuhan Hankou Hospital China, metadata accessed through the supplementary tables of the associated publication ^22^ and TCR-seq data obtained from the iReceptor platform ^43^ (http://ireceptor.irmacs.sfu.ca).

### Data pre-processing

All TCR-seq data was pre-processed for standardized analysis with Immunarch v0.6.6 ^44^. Data obtained from the Adaptive Biotechnologies ImmuneCODE database were used directly as inputs for Immunarch processing, with 1,475 COVID-19 patient samples and 88 healthy donor patient samples (1,563 samples total) successfully loaded and used for further analysis. For the ISB-Swedish cohort, patients were first filtered by those were sequenced by 10X Genomics. Sequence filtering and processing was performed as follows: for cells with multiple TRA and TRB CDR3 sequences, the first instance respectively were selected; only cells with paired TRA and TRB sequences were kept (column chain_pairing = Single pair, Extra alpha, Extra beta or Two chains); sequence files were converted to VDJtools format for input into Immunarch. COVID severity scores were translated from the WHO Ordinal Scale (0-7) to four tiers: healthy donor (0), mild (1-2), moderate (3-4), and severe (5-7). After pre-processing, the CD4 and CD8 datasets were composed of 136,429 and 69,687 clones, represented in a total of 16 healthy donors, 61 mild, 42 moderate, and 24 severe patients, 143 individuals total (16 healthy donors, 108 mild, 93 moderate, and 49 severe repertoires when accounting for patients with samples from two time points, 266 samples total). For the PLA General Hospital and Wuhan Hankou Hospital China cohort, cells with more than one TRA or TRB sequence had the chain with the highest number of reads kept for further analysis and sequence files were converted to VDJTools format for input into Immunarch. The PLA General Hospital aggregated patient dataset contained 31951 clones across 3 healthy donors (two healthy donors from the original study were excluded for lack of TCR CDR3 amino acid data), 7 moderate, 4 severe, and 6 convalescent patients (of which 4 were the second time point collections of moderate patients – P01, P02, P03, and P04). The Wuhan Hankou Hospital China aggregated patient dataset contained 42001 clones across 5 healthy donors, 5 moderate, and 5 severe patients. Metadata was manually reformatted from supplementary tables.

### Immune repertoire statistics

Clonotype statistics and diversity metrics were calculated using Immunarch v0.6.6 ^44^. For total number of unique clonotypes, the repExplore function was used with parameter .method = “volume”; for distribution of CDR3 sequence lengths, repExplore function with .method = “len” and .col = “aa”; for Chao1 estimator, repDiversity function with .method = “chao1”; for Gini-Simpson index, repDiversity function with .method = “gini.simp”; for Inverse Simpson index, repDiversity function with .method = “inv.simp”. Clonal proportion estimates were calculated with the repClonality function with .method = “top”. CDR3, V gene, and J gene usage proportions were calculated and aggregated directly from sample TCR data. Statistical significance testing comparing groups were performed using the two sided Wilcoxon rank-sum test by the wilcox.test in R.

### K-mer analyses

For K-mer abundance calculations, each VDJtools formatted sample was converted to a vector of CDR3 sequences. The vector was converted to k-mer statistics using the getKmers function from Immunarch, then merged with k-mer statistics of other samples using the R function merge with parameter all = TRUE for full outer join. Empty cells were converted from NAs to 0 counts. The 50,000 top variance unique k-mers were selected for downstream analyses (PCA and machine learning pipelines) with the exception of 3-mers which had 6916 unique k-mers. K-mer counts were normalized to sum to 1 for each sample prior to downstream analyses. PCA was performed using the prcomp function in R with parameter center = TRUE.

### Motif analyses

TCR clustering and specificity group analysis was performed using GLIPH2 ^29^. Software executable for analysis was obtained from http://50.255.35.37:8080/ and run with the human v2.0 reference on clonal data for each disease condition and T cell type. Parameters include global_convergence_cutoff=1, local_min_OVE=10, kmer_min_depth=3, simulation_depth=1000, p_depth=1000, ignored_end_length=3, cdr3_length_cutoff=8, motif_distance_cutoff=3, all_aa_interchangeable=1, kmer_sizes=2,3,4, and local_min_pvalue=0.001000.

Generation probability calculations were performed using OLGA ^30^. Software installation and setup was performed as described in https://github.com/statbiophys/OLGA and run on clonal data for each disease condition and T cell type. Representative calculation with parameters are as follows: olga-compute_pgen - i input.tsv --humanTRB -o out_pgens.tsv --v_in 1 --j_in 2.

### Single cell transcriptome analyses

Single cell transcriptome data from the ISB-S dataset were processed using Seurat v4.0.4. Pipeline included log normalization with scale factor 1,000,000, scaling and centering, PCA, nearest-neighbor graph construction, clustering with the Louvain algorithm, UMAP, differential gene expression, and generation of various visualizations. Parameters included: for the FindNeighbors function, dims = 1:10; for FindClusters, resolution = 0.6; for RunUMAP, dims = 1:10; for FindAllMarkers, only.pos = TRUE, min.pct = 0.25, logfc.threshold = 0.25. Differential gene expression between clonally expanded clusters and all other cells were performed using a downsampled cell subset (5,000 cells per group) of the data and the FindMarkers function with parameters logfc.threshold = 0.01 and min.pct = 0.1. P-value adjustment was performed using Bonferroni correction. Upregulated or downregulated genes with significance q-value < 1e-4 were then used for functional annotation with DAVID analysis. In addition to default Seurat outputs, custom R scripts were used to generate visualizations including UMAPs associated with CDR3 motifs and disease severity.

### Training and evaluation of machine learning models

Five ML-based approaches were trained on the k-mer frequency matrix generated from amino acids in the CDR3 region in the T cell repertoires of healthy donor and COVID-19 patients from the ISB-S datasets, using Python v3.8.6 and scikit-learn v0.23.1. These algorithms were: Random Forests (RF), Support Vector Machines (SVM), Naïve Bayes (NB), Gradient Boosting Classifiers (GBC) and K-Nearest Neighbors (KNN). The k-mer frequency matrix dataset was partitioned into subsets to perform binary classification between the healthy donor and the specified disease phenotype, such that models were trained for distinct classification tasks: healthy donor vs. mild disease, healthy donor vs. moderate disease, and healthy donor vs. severe disease. To address imbalanced datasets, healthy donor samples were upsampled to be equal to the number of COVID-19 samples represented in the dataset, prior to training. RFs were trained with 100 estimators, gini impurity criterion for measuring the quality of splits, minimum samples required to split an internal node of 2, minimum number of samples required to be a leaf node of 1, and bootstrapping to build trees. SVCs were trained with polynomial kernel and parameters C=20, degree=5, and probability=True. NBs were trained with default settings. GBCs were trained with 100 estimators, learning rate of 1.0 and maximum depth of 1. KNNs were trained with leaf size of 30 and the minkowski distance metric. Estimators were trained and evaluated with stratified 5-fold cross-validation, using 80% of the data for training and 20% of the data for validation, which was performed with 100 repetitions using the RepeatedStratifiedKFold function from sklearn. Plotly v5.1.0 was used to generate ROC plots from performance results.

### Statistical information summary

Comprehensive information on the statistical analyses used are included in various places, including the figures, figure legends and results, where the methods, significance, p-values and/or tails are described. All error bars have been defined in the figure legends or methods. Standard statistical calculations such as Spearman’s rho were performed in R with functions such as “cor”.

### Code availability

Key codes used for data analysis or generation of the figures related to this study has been included in this article and its supplementary information files, and have been deposited to GitHub at https://github.com/parkjj/tcrcov. Additional scripts used are also available upon request to the corresponding author.

### Data and resource availability

The authors are committed to freely share all COVID-19 related data, knowledge and resources to the community to facilitate the development of new treatment or prevention approaches against SARS-CoV-2 / COVID-19 as soon as possible. All relevant processed data generated during this study are included in this article and its supplementary information files or are currently being deposited into publicly accessible repositories. Raw data are from various sources as described above. All data and resources related to this study are freely available upon request to the corresponding author.

### Graphical illustrations

Certain graphical illustrations were made with BioRender (biorender.com).

## Supplemental Figure Legends

**Figure S1.**
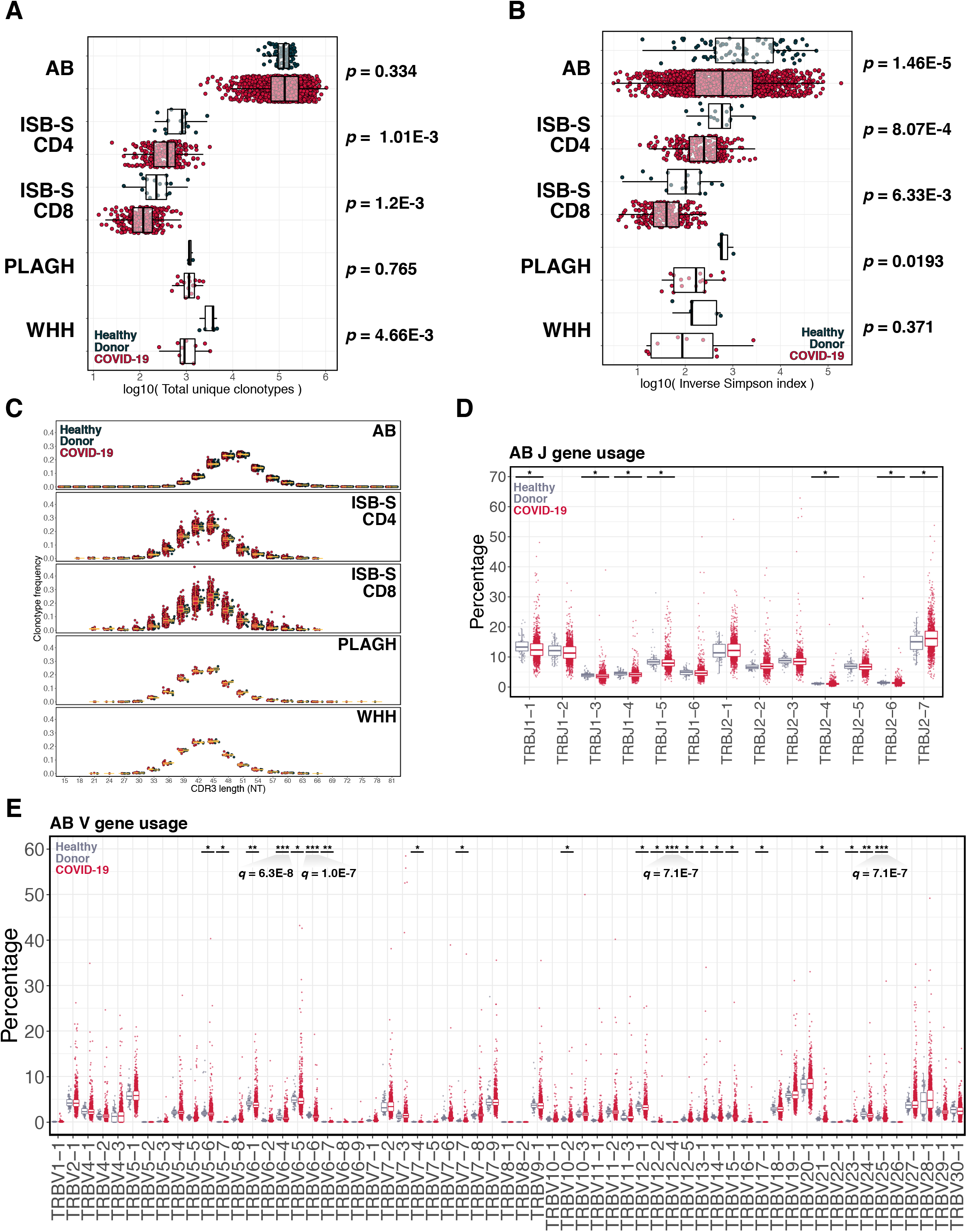
Diversity metrics and gene usages for TCR repertoire datasets. **(A)** Boxplot of total unique clonotypes for COVID-19 patients and healthy donors for each repertoire dataset. P-values were obtained using the two-sided Wilcoxon rank-sum test. **(B)** Boxplot of inverse Simpson indices for COVID-19 patients and healthy donors for each repertoire dataset. P-values were obtained using the two-sided Wilcoxon rank-sum test. **(C)** Boxplot of clonotype frequencies by CDR3 length for COVID-19 patients and healthy donors for each repertoire dataset. **(D)** Boxplot of J gene usages for samples from the AB dataset. Gray dots represent healthy donor samples; red dots represent COVID-19 samples. Statistical significance determined using the two-sided Wilcoxon rank-sum test and adjusted using the Benjamini & Hochberg method. * adj. P < 0.05, ** adj. P < 1e-4, *** adj. P < 1e-6. **(E)** Boxplot of V gene usages for samples from the AB dataset. Gray dots represent healthy donor samples; red dots represent COVID-19 samples. Statistical significance determined using the two-sided Wilcoxon rank-sum test and adjusted using the Benjamini & Hochberg method. * adj. P < 0.05, ** adj. P < 1e-4, *** adj. P < 1e-6.

**Figure S2.**
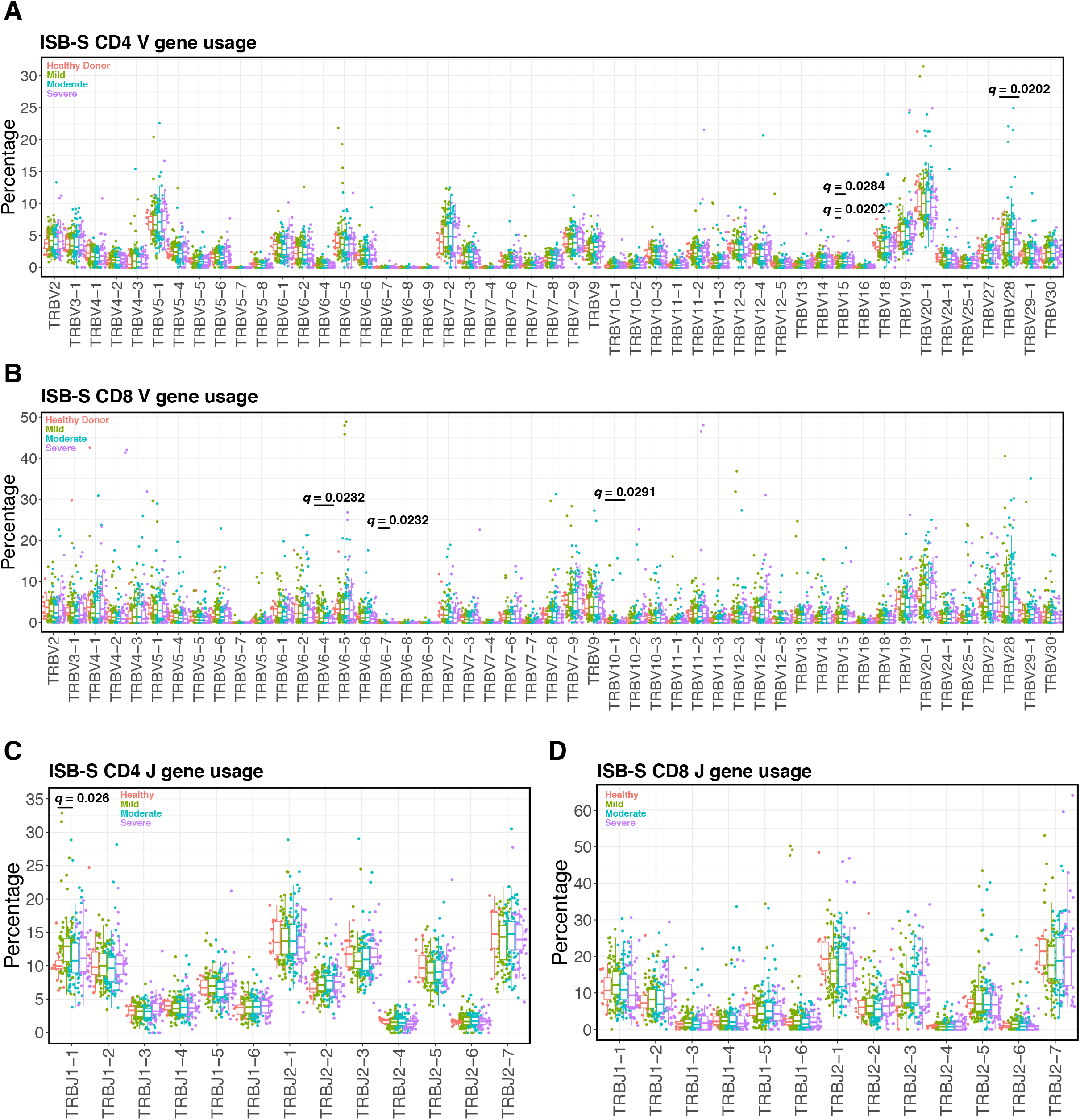
V and J gene usages by disease severity for ISB-S datasets. **(A)** Boxplot of V gene usages for CD4 samples from the ISB-S dataset. Red dots represent healthy donor samples; green dots represent mild; blue dots represent moderate; purples dots represent severe. Statistical significance determined using the two-sided Wilcoxon rank-sum test and adjusted using the Benjamini & Hochberg method. Adj. P < 0.05 labelled on plot. **(B)** Boxplot of V gene usages for CD8 samples from the ISB-S dataset. **(C)** Boxplot of J gene usages for CD4 samples from the ISB-S dataset. **(D)** Boxplot of J gene usages for CD8 samples from the ISB-S dataset.

**Figure S3.**
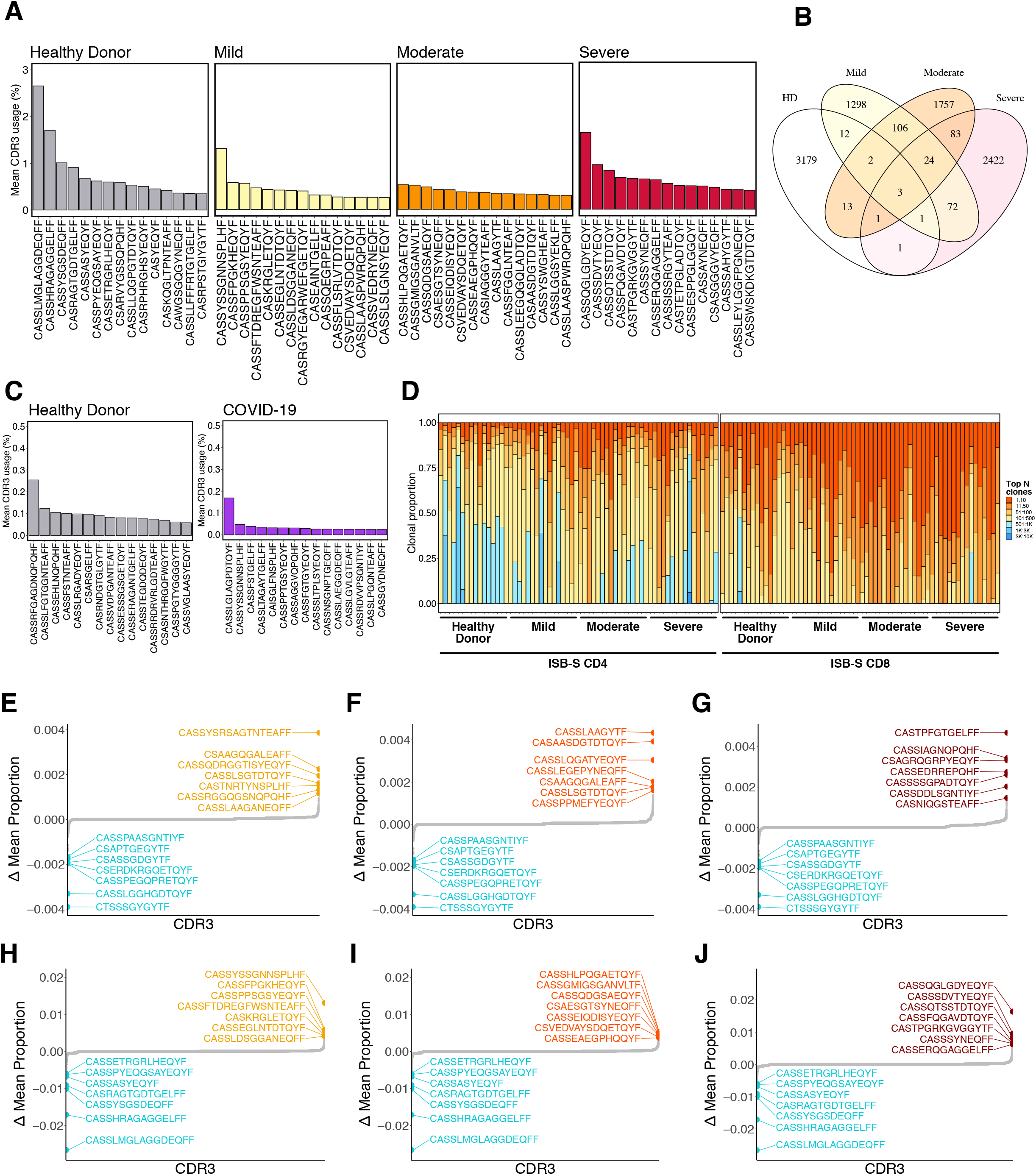
Additional CDR3 gene usage statistics. **(A)** Bar plots showing the top 15 mean CDR3 usages for patients in the ISB-S CD8 dataset grouped by disease severity (healthy donor = 16, mild = 108, moderate = 93, severe = 49). **(B)** Venn diagram showing overlap of top mean CDR3 usages (proportion threshold = 0.0001) for patients in the ISB-S CD8 dataset grouped by disease severity. **(C)** Bar plots showing the top 15 mean CDR3 usages for patients in the AB dataset grouped by disease status (healthy donor = 88, COVID-19 = 1,475). **(D)** Bar plot depicting relative abundance for groups of top clonotypes by disease condition for sampled repertoires (n = 16 per condition) from ISB-S datasets. **(E)** Dotted waterfall plot of CDR3 gene usage differentials between mild disease COVID-19 patients and healthy donors (delta mean proportion) in the ISB-S CD4 dataset. Yellow dots are CDR3 sequences enriched in moderate disease repertoires; light blue dots are CDR3 sequences enriched in healthy donors; grey dots are all other CDR3 sequences. **(F)** Dotted waterfall plot of CDR3 gene usage differentials between moderate disease COVID-19 patients and healthy donors in the ISB-S CD4 dataset. Orange dots are CDR3 sequences enriched in moderate disease repertoires. **(G)** Dotted waterfall plot of CDR3 gene usage differentials between severe disease COVID-19 patients and healthy donors in the ISB-S CD4 dataset. Red dots are CDR3 sequences enriched in severe disease repertoires. **(H)** Dotted waterfall plot of CDR3 gene usage differentials between mild disease COVID-19 patients and healthy donors in the ISB-S CD8 dataset. Yellow dots are CDR3 sequences enriched in mild disease repertoires. **(I)** Dotted waterfall plot of CDR3 gene usage differentials between moderate disease COVID-19 patients and healthy donors in the ISB-S CD8 dataset. Orange dots are CDR3 sequences enriched in moderate disease repertoires. **(J)** Dotted waterfall plot of CDR3 gene usage differentials between severe disease COVID-19 patients and healthy donors in the ISB-S CD8 dataset. Red dots are CDR3 sequences enriched in severe disease repertoires.

**Figure S4.**
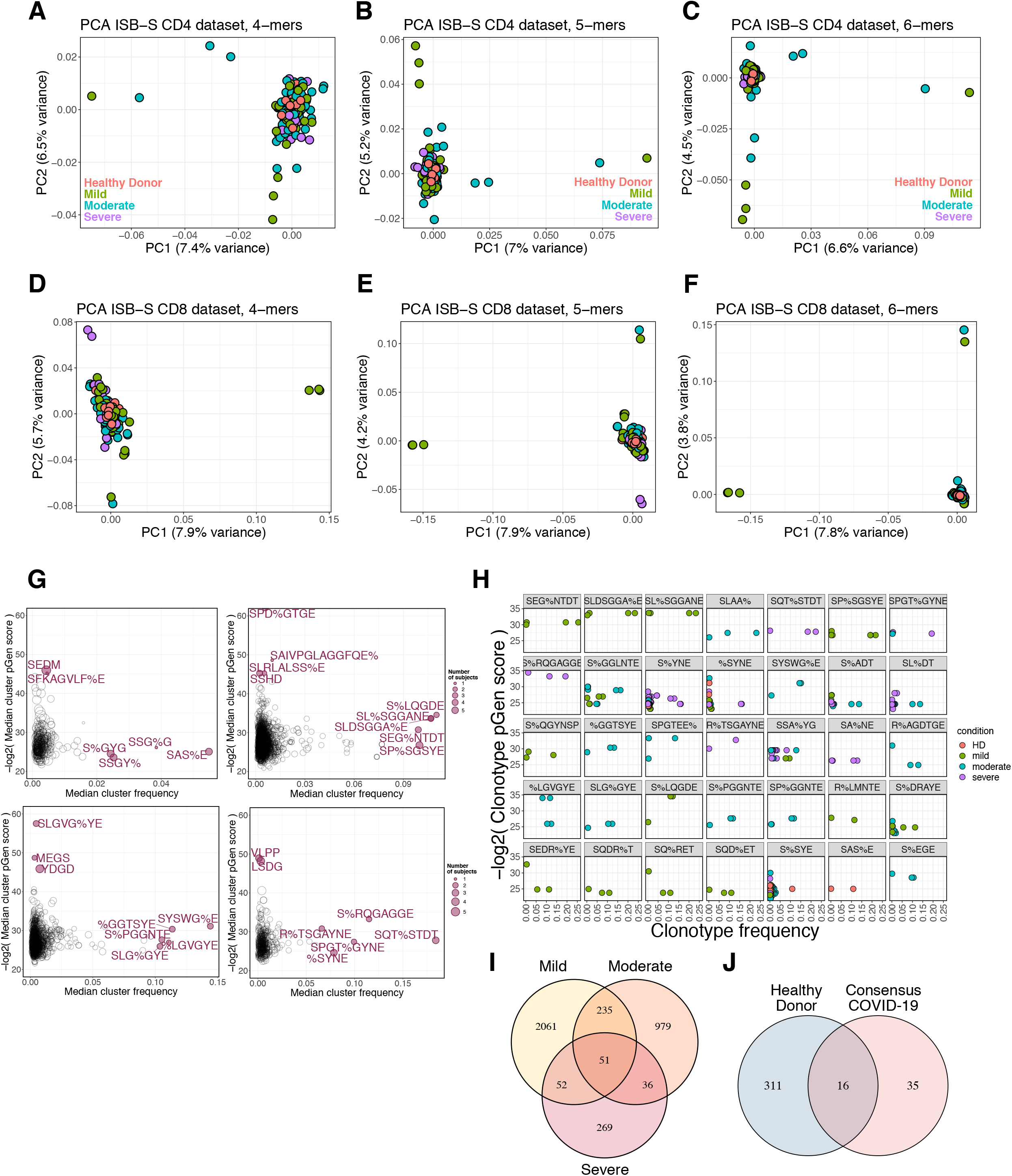
Additional k-mer and motif analyses. **(A)** PCA of 4-mer representations of TCR repertoires from the ISB-S CD4 dataset. **(B)** PCA of 5-mer representations of TCR repertoires from the ISB-S CD4 dataset. **(C)** PCA of 6-mer representations of TCR repertoires from the ISB-S CD4 dataset. **(D)** PCA of 4-mer representations of TCR repertoires from the ISB-S CD8 dataset. **(E)** PCA of 5-mer representations of TCR repertoires from the ISB-S CD8 dataset. **(F)** PCA of 6-mer representations of TCR repertoires from the ISB-S CD8 dataset. **(G)** Median frequency and pGen scores of COVID-19 and healthy donor associated T cell clusters from GLIPH2 analysis of the ISB-S CD8 dataset, grouped by disease condition. **(H)** Detailed view of frequencies and pGen scores of specific clonotypes associated with high frequency T cell clusters from CD8 dataset. Clonotypes are colored by patient disease condition. **(I)** Venn diagram showing overlap of COVID-19 associated T cell clusters for patients in the ISB-S CD8 dataset grouped by disease condition. **(J)** Venn diagram showing overlap between consensus COVID-19 associated T cell clusters (taken from intersection of disease conditions) and healthy donors for repertoires in the ISB-S CD8 dataset.

**Figure S5.**
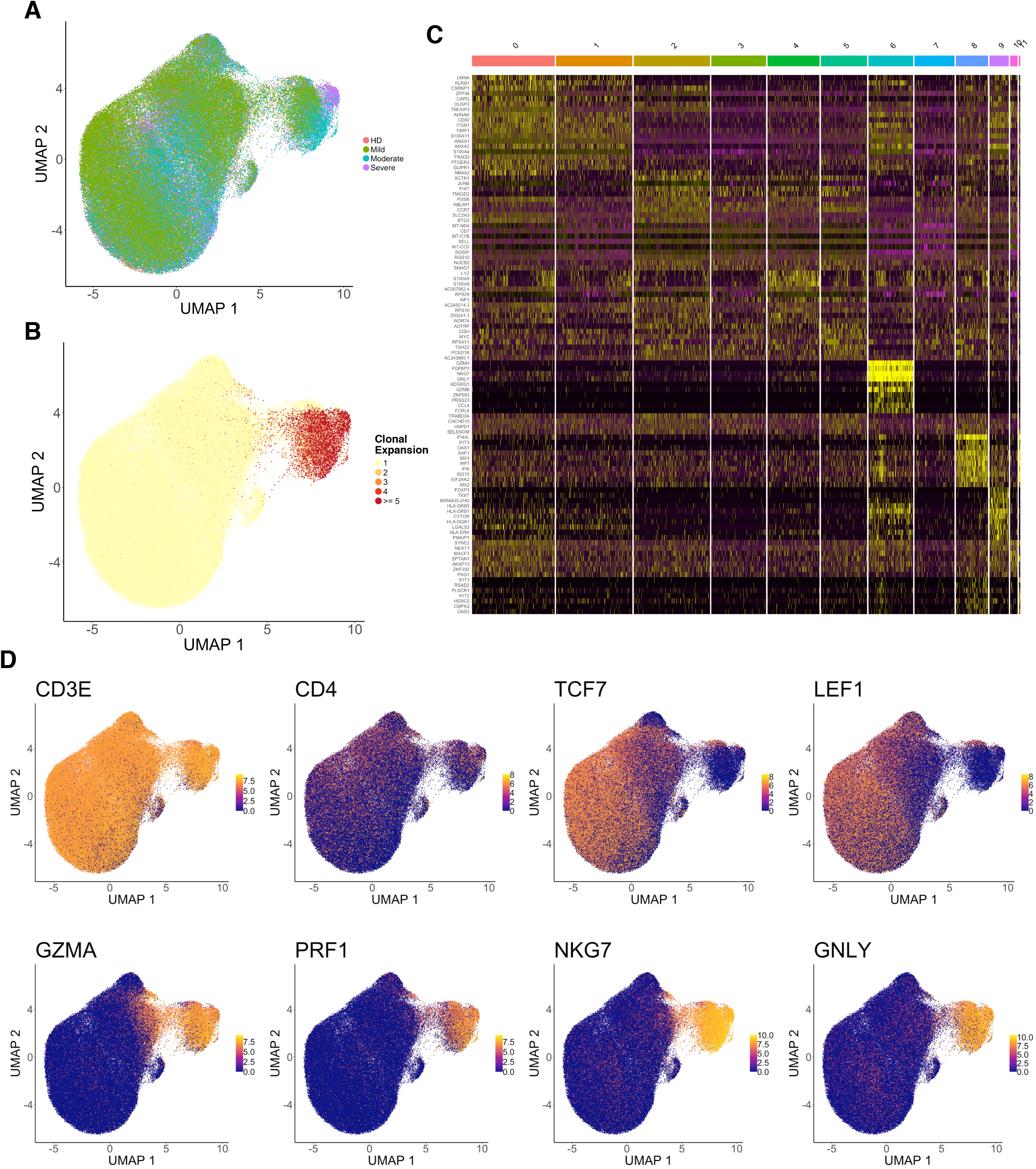
Additional single cell transcriptional analyses for CD4 T cells. **(A)** UMAP visualization of 137,075 CD4 T cell single cell transcriptomes from the ISB-S CD4 dataset labelled by disease condition. **(B)** UMAP visualization of CD4 T cell single cell transcriptomes labelled by clonal expansion. **(C)** Heatmap of differentially expressed markers for all identified clusters (n = 12). **(D)** UMAP visualizations highlighting expression levels of individual genes for cell phenotyping.

**Figure S6.**
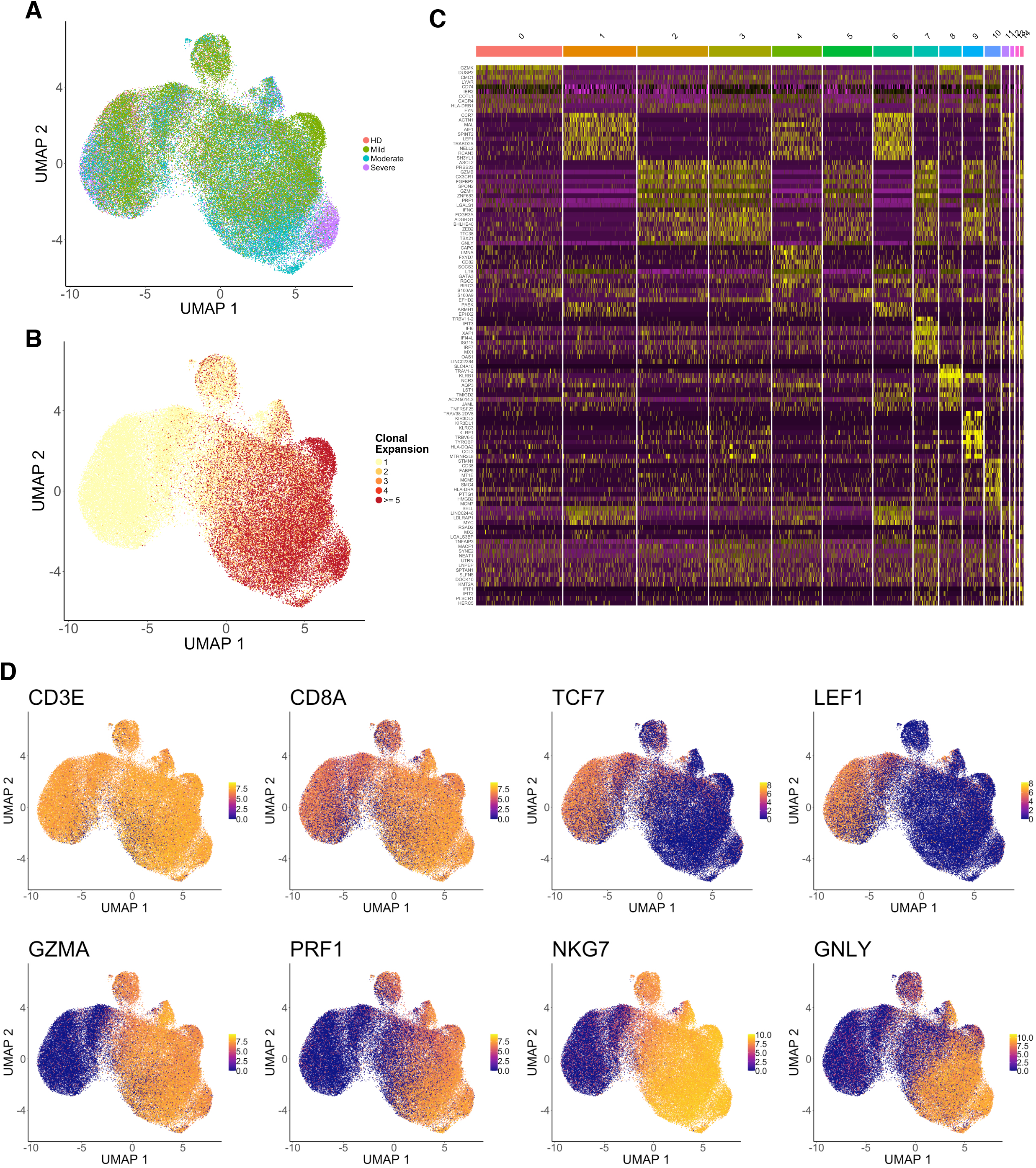
Additional single cell transcriptional analyses for CD8 T cells. **(A)** UMAP visualization of 70,237 CD8 T cell single cell transcriptomes from the ISB-S CD8 dataset labelled by disease condition. **(B)** UMAP visualization of CD8 T cell single cell transcriptomes labelled by clonal expansion. **(C)** Heatmap of differentially expressed markers for all identified clusters (n = 15). **(D)** UMAP visualizations highlighting expression levels of individual genes for cell phenotyping.

**Figure S7.**
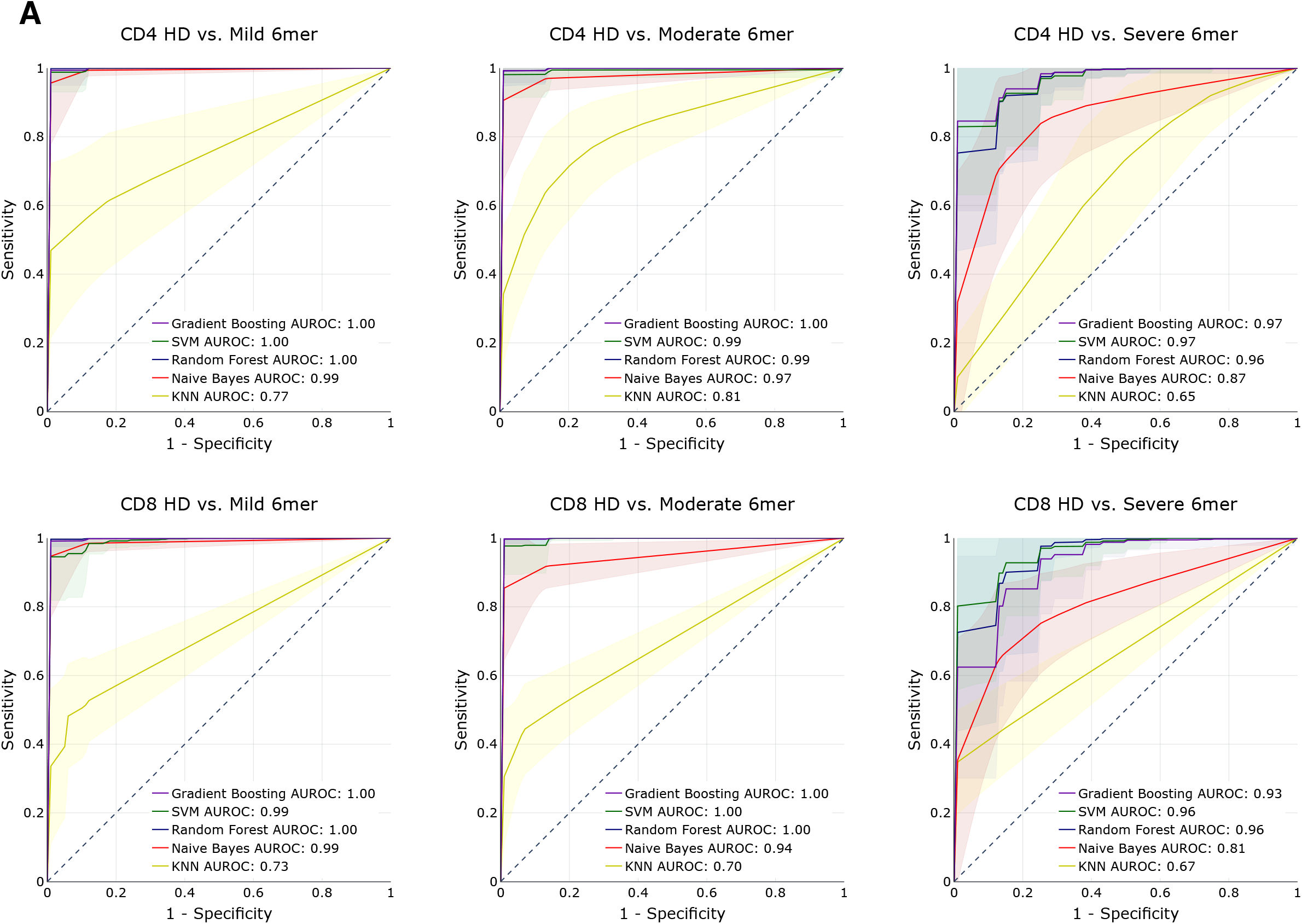
Additional machine learning analyses for disease severity. **(A)** AUROC curves for five machine learning models using 6-mer representations of TCR repertoire data. Models were trained to predict disease severity (mild, moderate, severe) vs healthy donors for CD4 (top row) and CD8 (bottom row) samples. Training and evaluation was performed using 100 repetitions of 5-fold cross-validations per model, average performance +/− 1 standard deviation shown on individual plots.

## Supplementary Datasets

### Supplementary Dataset

Dataset S1. Metadata for TCR repertoire samples obtained for all datasets used in study.

Dataset S2. Number of clones and unique clonotypes for each sample across datasets.

Dataset S3. CDR3 length statistics for each sample across datasets.

Dataset S4. Diversity statistics including Chao1 estimators, Gini-Simpson indices, and inverse Simpson indices for each sample across datasets.

Dataset S5. V and J gene usage statistics for each sample in Adaptive Biotechnologies datasets.

Dataset S6. V and J gene usage statistics for each sample in ISB-Swedish datasets.

Dataset S7. Principal components analysis results for 3-mer, 4-mer, 5-mer, and 6-mer representations of each sample in ISB-Swedish datasets.

Dataset S8. GLIPH clustering analysis patterns, scores, and statistics for ISB-Swedish datasets by T cell type and disease condition.

Dataset S9. OLGA analysis inputs of structured ISB-Swedish datasets by T cell type and disease condition.

Dataset S10. OLGA analysis output pGen scores of ISB-Swedish datasets by T cell type and disease condition.

Dataset S11. COVID-19 associated clusters in ISB-Swedish datasets by T cell type.

Dataset S12. UMAP coordinates for CD4 and CD8 T cell single cell transcriptome analyses.

Dataset S13. Cell proportions and counts for clonally expanded groups from CD4 and CD8 T cell single cell transcriptome analyses.

Dataset S14. Differential gene expression for Cluster 6 vs all other cells in CD4 T cell transcriptome analysis and Expanded group cells vs all other cells in CD8 T cell transcriptome analysis.

Dataset S15. Upregulated and downregulated genes for CD4 and CD8 T cell clonal expansion differential gene expression analysis using threshold q-value < 1e-4.

Dataset S16. DAVID gene ontology biological process annotations for CD4 and CD8 T cell clonal expansion differential gene expression analysis using threshold q-value < 1e-4.

Dataset S17. Average AUROC scores for machine learning models trained to predict disease severity from healthy donors using different k-mer representations of TCR repertoires.

